# Extracellular vesicles from human adipose stem cells are neuroprotective after stroke in rats

**DOI:** 10.1101/2020.11.18.388355

**Authors:** Francieli Rohden, Luciele Varaschini Teixeira, Luis Pedro Bernardi, Nicolly Paz Ferreira Marques, Mariana Colombo, Geciele Rodrigues Teixeira, Fernanda dos Santos de Oliveira, Elizabeth Obino Cirne Lima, Fátima Costa Rodrigues Guma, Diogo Onofre Souza

**Author notes:** Corresponding author: Diogo Onofre Souza; Postal address: Ramiro Barcelos, 2600, Laboratory 28.

## Abstract

Ischemic stroke is a prominent cause of death and disability, demanding innovative therapeutic strategies. Accordingly, extracellular vesicles (EVs) released from mesenchymal stem cells are promising tools for stroke treatment. In this study, we evaluated the potential neuroprotective properties of EVs released from human adipose tissue stem cells (hAT-MSC), which were obtained from a healthy individual submitted to liposuction. A single intranasal EVs administration was performed 24 h after the ischemic stroke in rats. The EVs brain penetration and the tropism to brain zone of ischemia was observed 18 h after administration. Thus, we measured EVs neuroprotection against the ischemic stroke-induced impairment on long-term motor and behavioral performance. Indeed, one single intranasal EVs administration reversed the stroke damages on: i) front paws symmetry; ii) working memory, short- and long-term memory; iii) anxiety-like behavior. These findings highlight hAT-MSC-derived EVs as a promising therapeutic strategy in stroke.

## INTRODUCTION

Stroke is a prominent cause of death and permanent disability worldwide, impacting health and financial burdens on society^1, 2^. Ischemic stroke most common symptoms are motor and cognitive impairment^3–5^. Currently it is known that ischemic stroke represents ~85% of stroke cases^6^, in which approximately 80% cause contralateral upper limb paresis^7^ and sensory function impairment^8^. In addition, clinical observations show that 20–50% of patients experience memory disorders^9, 10^.

The ischemic stroke pathophysiology characterizes by blood flow obstruction of a restricted brain region, forming an infarct nucleus, surrounded by an area known as penumbra zone^11^. Reperfusion of the penumbra zone contributes to a reduction in the final infarct size and reversal of neurological and behavioral deficits and consequently therapeutic strategies focus on this penumbra zone^12^. Currently, the gold standard treatment for ischemic stroke is the use of thrombolytic agents, which focuses on optimizing the reperfusion time in the penumbra zone^12^. However, this strategy has to be strictly applied within 4.5 h after the first symptoms^2^. In addition, reperfusion injury, such as hemorrhage, is the most dangerous collateral effect after thrombolysis^13^. Thus, the thrombolytic treatment is contraindicated for a certain class of patients at risk for bleeding (diabetes and hypertension) or with the formation of large blood clots^14, 15^.

In keeping with this, it is essential to find new therapeutic strategies for ischemic stroke. In vivo studies using a rat model of brain ischemia demonstrates that treatment with mesenchymal stem cells (MSCs), increases the therapeutic window up to 24 h after insult^16–18^. Though MSCs therapy seems very promising, it may cause some impacting damage, such as immune system rejection and the risk of developing tumor tissue^9, 19^. In this way, improvements in MSC therapy have been made to optimize its efficacy. It is currently thought that MSCs protective effects are due to the release of extracellular vesicles (EVs), pointing to an innovative perspective of the EVs use for stroke treatment. In fact, EVs are small double membrane vesicles (30-200nm), generated and release by many cell types, with physiologically role of a cell-to-cell communication^20, 21^.

In comparison to MSCs, EVs alone have low immunogenicity, absence of vascular obstructive effect, reduced risk of secondary microvascular thrombosis, and the ability to cross the blood-brain barrier^22, 23^. In vivo reports demonstrated that systemic administration of EVs from MSCs in rats, 24 h after ischemic stroke, increased angiogenesis, neurogenesis, neurite remodeling with a long-term recovery range^24–31^. However, systemically administered EVs may undergo metabolization before reaching the brain tissue^30–33^. In fact, by intravenous administration, EVs were detected after 24 h, in peripheral organs such as the lungs, liver and spleen^32^.

Thus, it seems crucial to focus on a straightforward and noninvasive manner to administer EVs. Current research presents promising results by intranasal administration of the EVs from MSCs for the treatment of brain disorders^34^. Moreover, Perets et al^35^ described the distribution of EVs after intranasal administration in vivo, showing that they quickly reach the brain damaged site.

In this study, we investigated the neuroprotection of an intranasal administration of EVs released from human adipose tissue mesenchymal stem cell (hAT-MSCs), in rat model of focal ischemic stroke. More specifically, we demonstrated that EVs administration, 24 h after stroke, fully reversed impairment on motor function, working memory, short- and long-term memory and anxiogenic-like behavior. Since hAT-MSCs can be easily obtained from the adipose tissue of healthy individuals, the administration of hAT-MSCs-derived EVs offers an accessible, affordable and effective therapeutic strategy for treating ischemic stroke

## MATERIALS AND METHODS

Human Adipose Tissue Mesenchymal Stem Cells (hAT-MSCs)

### Sources and culture of hAT-MSCs

The cells were from 2 sources:

a. Commercial hAT-MSCs: Cells were obtained from the POIETICS bank - Adipose-Derived Stem Cells (cat #PT-5006, donor 34464) were isolated from healthy human adults during elective surgical liposuction procedures. Cells were checked by the expression of CD13, CD29, CD44, CD73, CD90, CD105 and CD166, and absence of CD14, CD31 and CD45. They were tested negative for mycoplasma, bacteria, yeast and fungi. A frozen vial containing ~1 million cells was thawed at 37°C and plated in 25 cm^2^ flask (TPP). Cells were cultured with DMEM medium (Sigma) containing 10% FBS (Cripton), 100 units/mL penicillin (Gibco), 100 μg/mL streptomycin (Gibco), gentamicin 50 mg/L (Sigma) and fungizone 2.5 mg/L (Sigma). After 24 hours, the non-adherent and dead cells were removed. As adherent cells reached 80% confluence (passage 1: P1), hADSCs were detached with 0.25% trypsin/1mM ethylenediamine tetraacetic acid (EDTA) (Sigma) and plated in flasks at a density of 1,5×10^4^ cells/75 cm^2^ (passage 2: P2). The cellular density was determined by manually counting the number of cells at each division^36^. Identified as cell 1 (C1).
b. Patient-derived hAT-MSCs: cells were obtained from the subcutaneous fat of a 30-year-old female, submitted to abdominal liposuction at the Hospital de Clínicas in Porto Alegre, Brazil. The individual agreed to be part of the research and signed an informed consent form (GPPG 2018-0374). The fresh adipose tissue was washed with PBS buffer, minced and digested for 1 h in 0.1% collagenase at 37°C. The digestion process was stopped by the addition of Dulbecco’s Modified Eagle’s Medium (DMEM) containing 20% fetal bovine serum (FBS) and 100 units/mL penicillin (Gibco), 100 μg/mL streptomycin (Gibco). The digested suspension was filtered through a 70 μm nylon mesh cell filter to retain the tissue debris. The filtered suspension was centrifuged at 400×g for 5 min. The stromal vascular fraction (pellet) was resuspended in DMEM + 20% FBS (Cripton) medium and cultured in flasks cell culture of 25mc^2^ (TPP) at 37°C in a humidified atmosphere of 5% CO2. After 24 hours, the non-adherent cells were gently removed^36^. As adherent cells reached 80% confluence (passage 0: P0), confluent cells (hAT-MSCs) were detached with 0.25% trypsin/1mM ethylenediamine tetra acetic acid (EDTA) (Sigma) and plated in flasks at a density of 1,5×10^4^ cells/75 cm^2^ (TPP) (passage1: P1). Cells were cultured with DMEM medium (Sigma) containing 10% FBS (Cripton) and 100 units/mL penicillin (Gibco), 100 μg/mL streptomycin (Gibco), gentamicin 50mg/L (Sigma) and fungizone 2.5mg/L (Sigma). Identified as cell 2 (C2).

Both cells C1 and C2 were expanded under the same conditions and used only from 4^th^ to 8^th^ passage^26, 34, 37^.

### Cell Characterization

For characterizing hAT-MSCs, trypsinized cells were centrifuged (400×g for 5 min at room temperature), the cell pellet was resuspended in DMEM+10% FBS and the cells were counted in a Neubauer chamber. Shortly after, cells were incubated with antibodies at concentration of 1:50 for 4 h at 37°C. Afterwards, the cell suspension was centrifuged at 400×g for 5 min at room temperature, and the cells pellet was resuspended in 200 μl of PBS. Ten thousand events were analyzed on the flow cytometer (BD FACSCalibur™)^38^. C1 and C2, only in passage 4 (P4), were characterized as hADSCs by the expression of CDs by flow cytometry: CD34 (FITC Mouse Anti-human CD34 BD Pharmingen), CD45 (Human CD45 FITC Conjugate, Invitrogen), CD90 (PE Mouse Anti-Human CD90 BD Pharmingen) and CD105 (Human CD105 R-PE conjugate, Invitrogen).

An aliquot of 1×10^4^ hAT-MSCs was placed on a slide and analyzed by immunofluorescence. Cells were maintained in culture conditions for 72 h to adhere to the coverslip. Then, cells were incubated for 4 h at 37°C with the same antibodies used for cytometry: CD34, CD45, CD90 and CD105, at a ratio of 1:500. The negative control was prepared by incubating only the secondary antibodies: Alexa Fluor 555 (Invitrogen) and Alexa Fluor 488 (Invitrogen). Cells were washed with PBS (4 times) to remove excessive antibody, followed by fixation with PFA 4% for 2 h. Cells were washed again with PBS, and the coverslips fixed with fluoromount (Sigma) onto a histological slide for further analysis. Images were acquired using an 8-bit gray scale confocal laser scanning microscope (Olympus FV1000). Approximately 10–15 sections with 0.7 μm thick confocal were captured parallel to the coverslip (XY sections) using a ×20 objective (Olympus, U plan-super-apochromat, UPLSAPO 60X). Z-stack reconstruction and analysis were conducted using ImageJ software (http://rsb.info.nih.gov/ij/).

### Extracellular Vesicles (EVs)

#### EVs isolation and purification

As hAT-MSCs cell culture (from P4 to P8) reached 80% confluence, DMEM+10% FBS medium was exchanged for DMEM FBS-free (to avoid isolate vesicles contamination by FBS proteins). After 72 h of culture, the medium was collected for vesicle isolation and cells remained in culture. Of note, to recover from stress caused by FBS removal, in this step we supplemented cells with DMEM+10% FBS for 72 h^26^.

For EVs isolation we collected medium and centrifuge (3 times) at 4°C, varying the G-force and time (1^st^ - 400×g for 15 min, 2^nd^ 2000×g for 15 min and 3^rd^ 10,000×g for 30 min). We combined all supernatants and filtered through a 0.22 μm membrane. To finish the insulation, a centrifugation of 120,000×g at 4°C was performed for 2 h. The supernatant was then discarded, PBS was used to wash the pellet containing EVs, and centrifuged again under the same conditions. Finally, we resuspended the pellet in 100 μl of PBS and stored at −20°C^36^. Thus, EVs protein content was quantified by bicinchoninic acid (BCA) assay (Thermo Scientific Pierce™)^27^. The vesicles isolated from C1 and C2 cells are here termed EV1 and EV2, respectively.

#### EVs Characterization

EVs were characterized by flow cytometry, and membrane proteins CD63 and CD81 identified^39^. We first incubated EVs with magnetic beads (Thermo Fisher - Scientific - Invitrogen™) coated with primary antibody CD63 (Exosome-Human CD63, Thermo Fisher - Scientific - Invitrogen™) and CD81 (Exosome-Human CD81, Thermo Fisher - Scientific - Invitrogen™) for 18 h at 4°C and stirring. For each preparation 10 μl of EVs suspension at 1 mg/ml was applied. After washing with PBS (3 times), CD63 (CD63 Mouse anti-Human, FITC, Clone: MEM-259, Invitrogen™) and CD81 antibodies (with no beads) were added to the solution containing the EVs (PE Mouse Anti-Human CD81 Clone JS-81, BD Pharmingen™). After 1 h of incubation, EVs were gently washed with PBS (to remove excessive antibody) and resuspended in 200 μl of PBS for analysis. Ten thousand events were analyzed by flow cytometry. Thus, we measure mean particle size and polydispersity index (PDI) using photon correlation spectroscopy. The hAT-MSCs-derived EVs suspension (50 μl), at concentration 1 mg/ml, were diluted in 1 ml of PBS. All analyses were carried out in triplicate using a Malvern Nano-ZS90® (Malvern Instruments, England) at 25°C.

#### EVs Purity Measurement

We applied transmission electron microscopy (TEM) analysis, using direct examination technique, to evaluate EV purity and to check diameter size^34^. EVs suspension (10 μL), at 1mg/ml of protein, was aliquoted onto a grid covered with carbon film (formvar/carbon) and dried at room temperature. Uranyl was used as a contrast. The sample was analyzed by TEM 120Kv (JEM 1200 Exll-JEOL).

#### EVs Labeling

EVs were labeled with red fluorescent membrane dye PKH26 (MINI26, Sigma). In brief, the EVs-containing PBS solution was centrifuged at 120,000×g for 2 h, at 4°C and resuspended with the diluent of the fluorescent kit. Filtered PKH26 (4mM) and EVs (200 μg/ml) were mixed at a ratio 1:1 for 5 min, followed by the addition of 5% BSA. EVs were washed with PBS and centrifuged under the same conditions. EVs pellet was resuspended in 0.5 mL of PBS. To eliminate any dye aggregates, the solution was filtered through a 0.2 μm membrane filter^34^.

### Animals

Adult (90–120 days old) male Wistar rats weighting 350–400g were maintained under controlled light (12/12 h light/dark cycle), 22°C ±2 with water and food *ad libitum.* All procedures were performed according to the Guide for the Care and Use of Laboratory Animals and the Brazilian Society for Neuroscience and Behavior (SBNeC), recommendations for animal care. The Ethics Committee for the Use of Animals at the Universidade Federal do Rio Grande do Sul (process number: 31888) approved this project.

#### Permanent Focal Ischemia Surgery

Anesthetized animals (ketamine hydrochloride - 90mg/kg, 450μl/kg i.p. and xylazine hydrochloride - 10mg/kg, 300μl/kg i.p.) were placed in the stereotaxic apparatus. After skin incision, the skull was exposed, and a craniotomy was performed by exposing the left frontoparietal cortex (+2mm to −6mm A.P. and −2mm to −4mm M.L. from the bregma). An ischemic lesion was induced by thermo-coagulation of motor and sensorimotor pial vessels^40^. Blood vessels were thermo-coagulated by placing a hot probe near the dura mater for 2 min, until red-brown color indicated complete thermo-coagulation. Soon after, the skin was sutured, and animals placed on a heating pad at 37°C, until full recovery from anesthesia. The animals were divided in: Naive; naive treated with EV (Naive + EV1 or EV2); ischemic animals treated with PBS (ISC); ischemic animals treated with 200 ug/kg EVs (ISC + EV1 or EV2). Animals were randomly allocated to different treatment groups.

#### EVs Intra Nasal Treatment

EVs intranasal treatment was performed 24 h after ischemic insult. Sedated animals (O2 flow rate of 0.8-1.0 L/min with Isoflurane levels of 2.5-3.0%), were slowly injected into the nasal cavity a single dose of 50 μL EVs (ISC+EVs) or 50 μL PBS (ISC). The EVs dose (200 μg/kg) was selected after performing a dose curve: ISC+PBS; ISC+100 μg/kg; ISC+200 μg/kg and ISC+300 μg/kg (n=3).

#### Extracellular Vesicles Detection (EVs) in Rat Brain

Distribution of EVs in rat brains was analyzed 18 h after intranasal administration of fluorescent EVs (PKH26-mini, Sigma)^31, 34, 35^. Anesthetized animals were (ketamine hydrochloride −90mg/kg, 450μl/kg i.p. and xylazine hydrochloride 10mg/kg, 300μl/kg i.p.) transcardially perfused using a peristaltic pump (10 mL/min, with PBS and followed by PFA 4% – 100mL of each). Brains were dissected and immersed in PFA 4%, pH 7.4 and stored for a maximum of 7 days at 4°C. Coronal brain sections 20 μm thick were obtained using a vibratome (Leica) +2.20mm from the rostral and −0.80mm caudally relative to bregma. Brain slices were mounted on glass slides and incubated for 5 min in the dark with 1μg/mL Hoechst dye (33342 Sigma-Aldrich) in PBS to detect cell nuclei. The slices were washed with PBS (4 times) and the slides fixed with fluoromount (Sigma).

Images were acquired using an 8-bit gray scale confocal laser scanning microscope (Olympus FV1000). Approximately 10-15 sections with 0.7 μm thick confocal were captured parallel to the coverslip (XY sections) using a ×60 objective (Olympus, U plan-super-apochromat, UPLSAPO 60X). Z-stack reconstruction and analysis were conducted using ImageJ software (http://rsb.info.nih.gov/ij/). Images were obtained from 3 brains of ISC + EV1 animals and 3 brains of naive animals. Six images of each brain were selected (3 contralateral and 3 ipsilateral of ISC + EV1, 3 left side and 3 right side of naive), totaling 72 images. To quantify the EV fluorescence, 15 random regions of interest were analyzed in the same place for each image to ensure a reliable quantification.

#### Cylinder Task (CT)

The cylinder task allows for evaluation of motor sequelae caused by ischemic insult^41^. We thus determinate animal motor symmetry of front paws. Exploration of the apparatus is considered only when rats raise their bodies and contact their front members on the cylinder wall (20 movements are counted). The apparatus consisted of a transparent glass cylinder 20 cm in diameter and 30 cm in height. All animals were submitted to this task, 2 h before surgery, to verify basal forelimb symmetry. The CT was repeated on the 3^rd^, 7^th^, 14^th^, 21^st^, 28^th^, 35^th^ and 42^nd^ day after EVs treatment. The performance was recorded using ANY-Maze software (Stoelting CO., Wood Dale, IL), and the ipsilateral (to the lesion), contralateral or both forelimbs preference were counted using a blind analysis. The asymmetry of each animal was calculated by the following formula: asymmetry = (% of ipsilateral paw use) - (% of contralateral paw use). The asymmetry percentage was converted into a symmetry percentage^42^. Groups: Naive (n=6), Naive+EV1 (n=6), ISC (n=22), ISC+EV1 (n=17); B) Naive (n=6), Naive+EV2 (n=6), ISC (n=22), ISC+EV2 (n=17). At the end of each task, the apparatus was cleaned using 70% ethanol solution.

#### Open Field Task (OFT)

The open field task evaluates habituation to novelty (assessing short- and long-term memory - exploratory activity) and locomotor activity^43^. The apparatus consisted of a black cage measuring 50 cm in length × 50 cm in width × 50 cm in height. The animals were placed individually for 10 min. Animals performed the task on the 7^th^, 21^st^ and 42^nd^ days after EVs treatment. Short-term memory (habituation to novelty) was evaluated considering the decrease of locomotion in the 0-5 min of the first session (7^th^ day). Long-term memory was considered as a decrease in locomotion in the 1^st^ min in the following sessions (from the 1^st^ to the 3^rd^ session). Groups: Naive (n=8), Naive+EV1 (n=7), ISC (n=6), ISC+EV1 (n=9) and ISC+EV2 (n=5).

At the end of each session, the apparatus was cleaned with 70% ethanol solution. The task was recorded and analyzed by ANY-maze 6.1 software.

#### Novel Object Recognition Task (NORT)

To acclimate the animals in the arena, we performed a 10 min OFT. We evaluated the rats short- and long-term memory^44^ by applying the NORT. The NORT was performed on the 7^th^, 21^st^ and 42^nd^ days after EVs treatment. After 90 min of interval from the end of the OFT, it was performed the NORT training session. The animals were individually placed on the periphery of the arena for exploration. Two identical (familiar) objects (FO1 and FO2) were placed in the arena, animals were allowed to explore for 10 min. Sniffing and touching the object were considered as exploratory behavior. To evaluate short-term memory, 90 min after the training session, each animal was placed back in the arena. One of the two objects used in the training session (FO) was replaced by a new object (NO). The long-term memory was evaluated 24 h after the short-term memory task. The animals were placed back in the arena with same FO used in the first test. For this test, we placed the NO in a different position. In all sessions, the time spent exploring the objects was recorded for 10 min, using ANY-maze 6.1 software. Results were expressed as a percentage of time exploring each object during the training/test session. Animals that recognize the familiar object as such (learn the task) explore the novel object more than 50% of the total exploring time. Groups: Naive (n=8), Naive+EV1 (n=7), ISC (n=6), ISC+EV1 (n=9) and ISC+EV2 (n=5). At the end of each task, the apparatus was cleaned using 70% ethanol solution.

#### Y-maze Task

To evaluate working memory, we applied a spontaneous alternation task in a Y-maze^45, 46^. The maze consisted of grey wooden walls with 3 identical arms (30 cm × 8 cm × 15 cm, at a 120° angle from the other arms). The Y-maze task assesses spontaneous alternation and was conducted as follows^43^. A decrease in the number of spontaneous alternations indicates impaired working memory. Animals freely explore the maze for 5 min, starting from the end of the first arm. Alternation was defined as a complete cycle of consecutive entries in each of the 3 arms. The percentage of alternation (PA) was calculated by the PA = number of alternations/total number of entries in each arm - 2) × 100^4748^. Groups: Naive (n=8), Naive+EV1 (n=4), ISC (n=7), ISC+EV1 (n=9) and ISC+EV2 (n=5). At the end of each task, the apparatus was cleaned using 70% ethanol solution. The parameters were recorded and analyzed using ANY-Maze 6.1 software.

#### Elevated Plus Maze Task (EPMT)

The EPMT evaluates anxiety-like behavior^48^. The apparatus consists of 2 enclosed arms (50 cm × 10 cm × 40 cm) divided perpendicularly by 2 open arms (50 cm × 10 cm). The apparatus was placed 70 cm above the floor and the EPMT performed for 5 min. The parameters were recorded and analyzed using ANY-Maze 6.1 software. An increase in the time spent in the open arms indicates anxiolytic-like behavior. The animals were evaluated only once, on the 7^th^ day post EV treatment. Groups: Naive (n=6), Naive+EV1 (n=6), ISC (n=11), ISC+EV1 (n=16) and ISC+EV2 (n=5). At the end of each task, the apparatus was cleaned using 70% ethanol solution.

### Statistical Analysis

One-way ANOVA was applied to analyze the fluorescent EVs in the brain. Twoway RM ANOVA was applied for CT, followed by Sidak’s multiple comparisons test. For the long-term memory test, we applied a two-way ANOVA followed by Sidak’s multiple comparisons test. Short-term memory was evaluated using an unpaired t-test. The Y-maze task was evaluated using 1-way ANOVA in a post-hoc Turkey’s test^49^. EPMT was analyzed with unpaired t-tests. In addition, unpaired T-tests were used for the NORT with a theoretical average of 50%. Data are reported as the mean ±SD. All analyses were performed using Graph Pad Prism 6.0.

## RESULTS

### Cells and EVs

#### Cell Characterization

Supplementary Figure 1. There is no single marker to characterize hADSCs, which is done by immunophenotyping based on the expression (>70%) of CD90 and CD105 associated with the absence of expression (<5%) of CD34 and CD45^26^. Displacement of the fluorescence peaking to the right side registers a positive value for protein markers. More than 70% of C1 and C2 cells were labeled for CD90 and CD105. Only 0.34% of the analyzed cells, C1 and C2, were marked for DC45 and CD34, which was a characteristic of these cells. There was no displacement of the fluorescence peaks. The cells were also identified by fluorescence microscopy.

#### Extracellular Vesicles Characterization

Figure 1. hADSC-derived EVs were isolated by differential centrifugation and the resulting pellet of enriched EVs was analyzed by flow cytometry, TEM and Zetasizer. By flow cytometry, EV1 and EV2 released from hADSCs C1 and C2, respectively, were observed. **A)**, **B)**, **C)** and **D)** show the fluorescence of EVs incubated only with beads (gray peak). In **A)** and **B)** the displacement of fluorescence peaks to the right, indicating a positive for CD63 (green). In **C)** and **D)**, the displacement of fluorescence peaks to the right, indicating a positive for CD81 (red). Thus, flow cytometry showed that more than 90% of the EVs released expressed CD63 and CD81, establishing EV characterization^39^.

**Figure 1.**
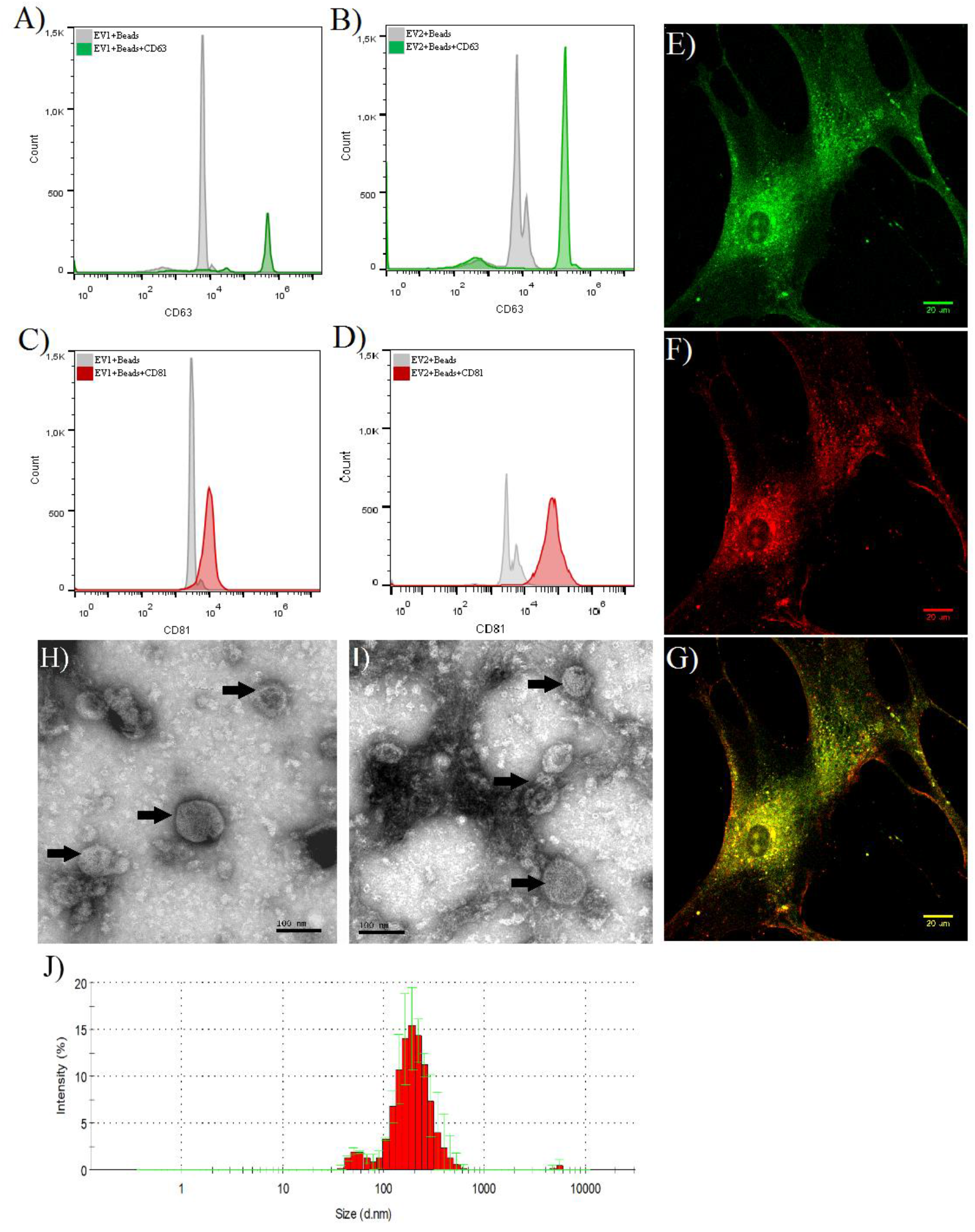
Representative images of EVs analysis by flow cytometry and fluorescence with CD labeling specific for EVs. The positive result (displacement of the fluorescence peak) for CD63 is labeled as green in **A**) EV1 and **B)** EV2. The positive result for **C)** EV1 and **D)** EV2 for CD81 is labeled as red. These results were compared to EVs incubated with BEADS only (grey). The fluorescence signal from these markers was observed inside C1 cells, CD63 (green) **(E)**, CD81 (red) (**F**), Merge **(G)**; fluorescence images were from confocal microscope, using the 60x objective. Transmission electron microscopy (TEM) images by direct examination, with 200k magnification, showing derived-hAT-MSC EVs purity and size, averaging 150nm is shown in C1 (**H**) and C2 (**I**). **(J)** Representative graph of the 150nm average diameter analysis done by Zetasizer instrument. N=3. Scale bars: 20μm.

In **E)** and **F)**, the purity of EV1 (E) and EV2 (F) isolation was confirmed by the TEM-direct technique. The suspension only consisted of released vesicles with cylindrical morphology and electron dense membranes, which were observed in both EV1 and EV2 (indicated by black arrow). The size of EVs was analyzed with a Zetasizer instrument (G), which confirmed the average diameter to be 150nm, with a polydispersity index (PDI) average of 0.3, giving evidence that most of our EVS had the same size. Through cell immunofluorescence, EVs inside hAT-MSCs C1 were observed. **H)**, **I)** and **J)** were positively marked for CD63, CD81 and merge, respectively. CD63 and CD81 were distributed in the plasma membrane and there is also a large indication near the nucleus.

#### Dose curve of EV1 administration on motor neuroprotective effect in ISC animals

Supplementary Figure 2. The cylinder task (CT), which measures front paw symmetry, was used for determining the lowest dose to be used for EV stroke treatment. On day 0 (24 h before stroke) all animals were evaluated by CT and only those presenting nearly 100% front paw symmetry were included in the study. On day 3 (2 days after stroke), all ISC animals presented a mean symmetry of 30%. EV1, or vehicle was intranasally administered 24 h after stroke. On the 42^nd^ day after intranasal administration, animals receiving vehicle, 100μg/kg, 200μg/kg or 300μg/kg presented symmetry recovery of 58%±6, 67%±3, 87%±8 and 82%±3, respectively. Animals receiving 200μg/kg or 300 μg/kg did not differ from naive group animals - 95%±3, pointing to a total recovery of symmetry. Thus, the dose of 200μg/kg at 24 h after stroke was selected. for the further experiments

#### Detection of Extracellular Vesicles in the brain

Figure 2. The distribution of EV1 (200μg/kg) in the brain was analyzed by fluorescence quantification (red): **A)** ISC ipsilateral penumbra zone; **B)** ISC contralateral side **C)** naive group animals, equivalent to the ipsilateral penumbra zone. The images indicate that EVs had tropism to the ISC ipsilateral penumbra zone. D)Figure representing the location of image acquisition; E) shows significant increase in fluorescence intensity in the ipsilateral penumbra zone compared to the contralateral side and naive group animals, thus indicating an EV1 tropism near the ipsilateral penumbra zone.

**Figure 2.**
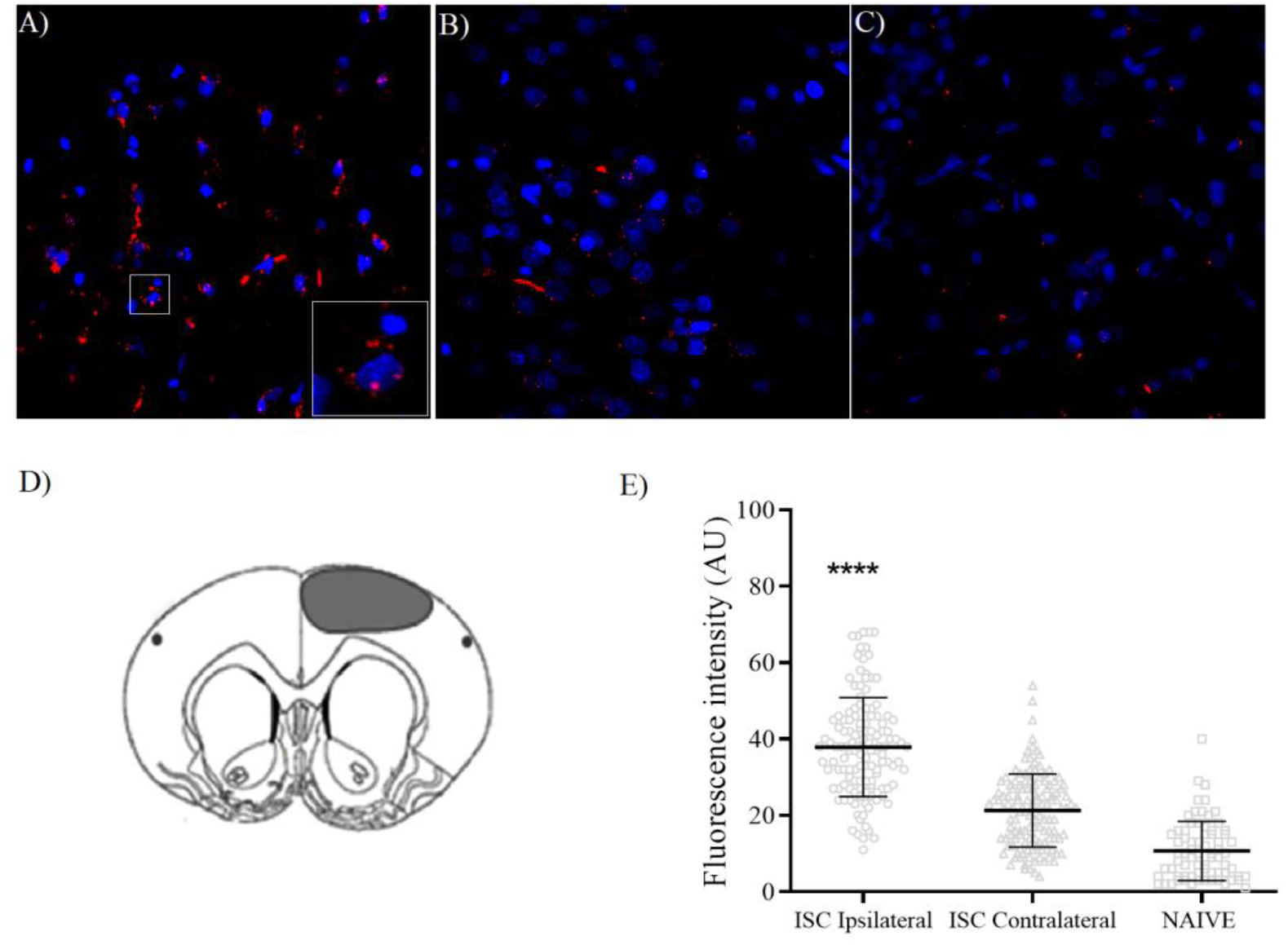
EVs distribution in brain cortex 18 hours after stroke. Representative images: **A)** penumbra zone in ISC+EV1 animal; **B)** contralateral side in ISC+EV1 animal; **C**) naive animal; **D)** representation of penumbra zone in brain ipsilateral region; **E)** mean brain fluorescence intensity was performed using ImageJ software. Statistical analysis by Ordinary one-way ANOVA. Data are reported as the mean ± S.D. [F (2,325) =144.0, p<0.0001]; ****p < 0.0001, comparing to all other groups. N=3. Scale bars: 20μm.

#### Effect of treatment with 200μg/kg of Extracellular Vesicles on neuromotor recovery

Figure 3. EV1 and EV2 treatment at 24 h after stroke caused a time dependent recovery of front paw symmetry from the 7^th^ day after treatment and attained total recovery from the 21^st^ day (EV2) or from the 28^th^ day (EV1).

**Figure 3.**
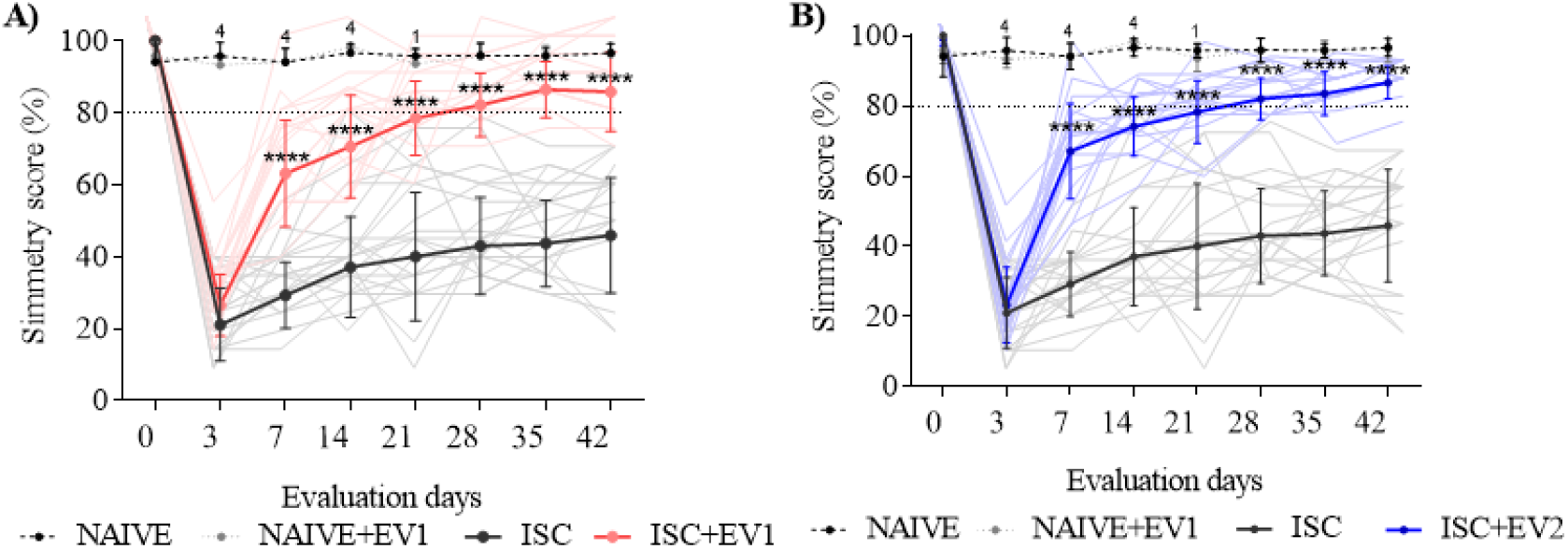
Symmetry score. The results are represented as group mean (dotted line and strong lines) and individual performances (soft lines). **A)** Naïve (n=6), Naive+EV1 (n=6), ISC (n=22), ISC+EV1 (n=17); **B)** Naïve (n=6), Naive+EV2 (n=6), ISC (n=22), ISC+EV2 (n=17). Data are expressed as mean ± SD, analyzed by two-way ANOVA followed by Tukey’s multiple comparisons test; ****p < 0.0001, compared to ISC group; ^1^p<0.05 and ^4^p<0.0001 compared to ISC+EV groups.

#### Open Field Task (OFT)

Figure 4. This task evaluates the locomotion of the animals in 3 sequential OFT exhibitions: 7^th^, 21^st^ and 42^nd^ days after EV1 or EV2 treatment. All groups presented short-term memory in the 1^st^ exhibition. Naive and Naive+EV1 groups also presented long-term memory. ISC animals presented long-term memory impairment, an effect reversed with EV1 and EV2 treatment.

**Figure 4.**
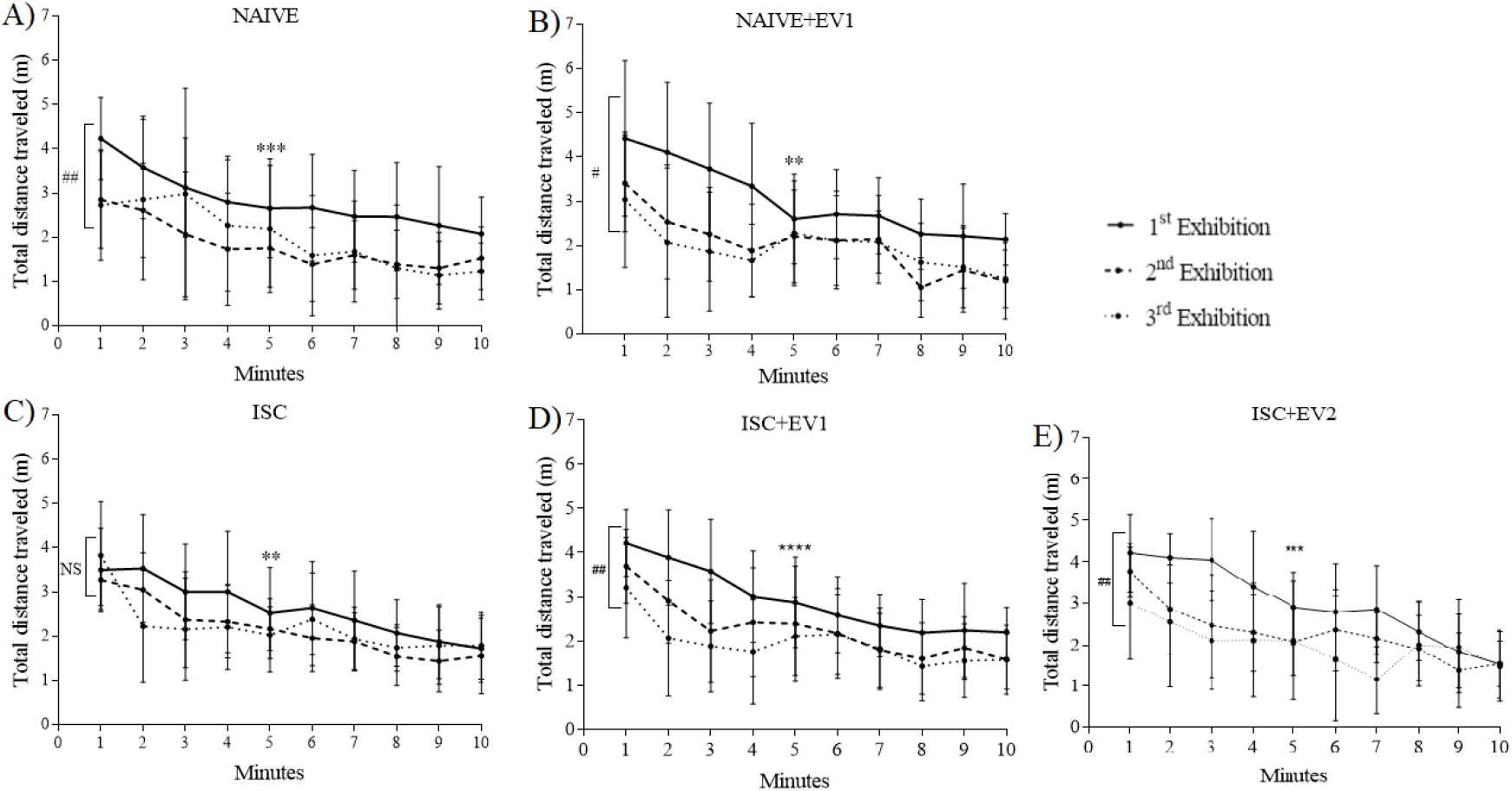
Open Field Task. Groups: Naive (n=8), Naive+EV1 (n=7), ISC (n=6), ISC+EV1 (n=9) and ISC+EV2 (n=5). Statistical analysis by 2way ANOVA, followed by Tukey’s multiple comparisons test. Data are reported as mean ± SD. **p < 0.01, ***p < 0.001, compared to the 1^st^ minute of the same session; #p<0.05, ##p<0.001, compared the 1^st^ minute of the first session with the 1^st^ minute of the second and third sessions.

#### Novel Object Recognition Task (NORT)

Figure 5. The NORT was used for evaluating short- and long-term memory on the 7^th^, 21^st^ and 42^nd^ days after EV1 or EV2 treatment. Stroke impaired the short-term memory (evaluated 90 min after the training session) and also the long-term memory (evaluated 24 h after the training session). EV1 or EV2 treatment reversed these stroke effects.

**Figure 5.**
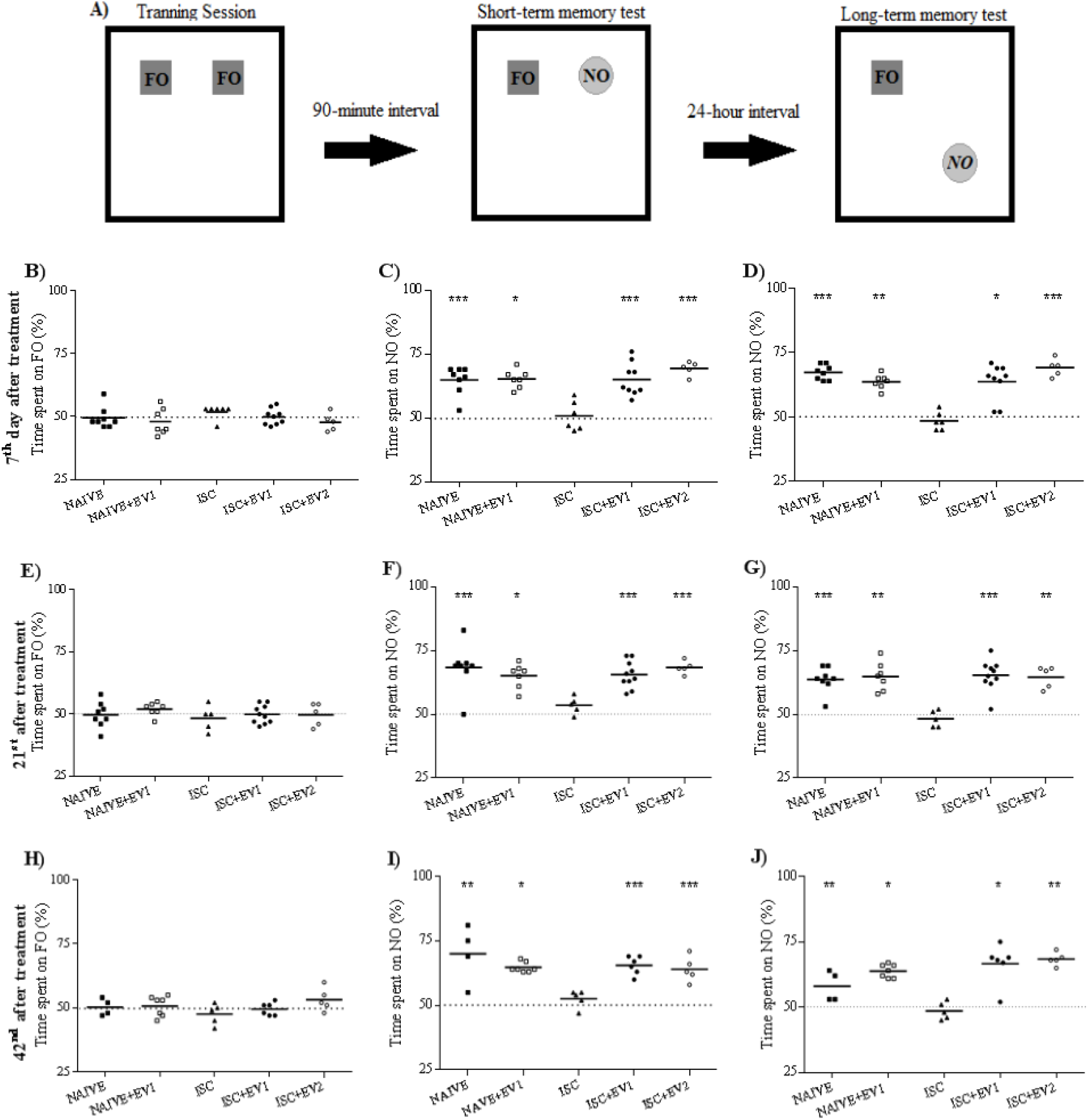
**A)** Scheme of protocol by using Familiar Object (FO) and/or Novel Object (NO). Groups: Naive (n=8), Naive+EV1 (n=7), ISC (n=6), ISC+EV1 (n=9) and ISC+EV2 (n=5). Training sessions using 2 FOs; STM test sessions 90 minutes later and LTM test sessions 24 hours later. **B**, **E** and **H)** 3 successive training sessions; **C**, **F** and **I)** 3 successive test STM sessions; **D**, **G** and **J)** 3 successive test LTM sessions. Unpaired T-test with theoretical average of 50%. Data reported as mean (SD less than 16% of each respective mean); *p < 0.05, **p < 0.01 and ***p < 0.001, comparing to theoretical average of 50%.

#### Y-maze Task

Figure 6. Spontaneous alternation in a Y-maze task is widely used to evaluate working memory. The task was performed on the 7^th^ day after treatment with EVs. Animals submitted to stroke presented working memory impairment, an effect reversed by EVs.

**Figure 6.**
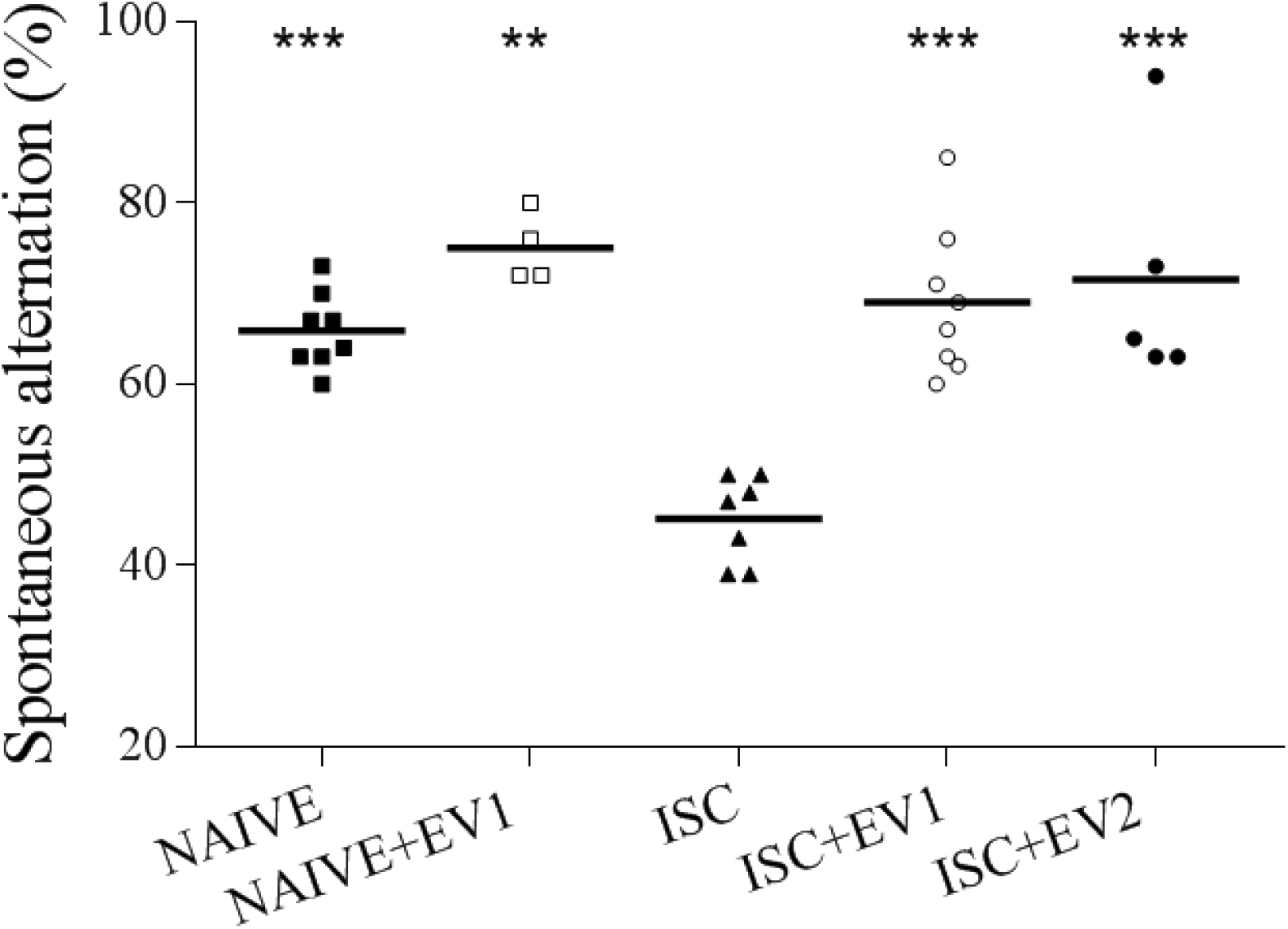
Y-maze task. Groups: Naive (n=8), Naive+EV1 (n=4), ISC (n=7), ISC+EV1 (n=9) and ISC+EV2 (n=5). 1-way ANOVA of a post-hoc Turkey’s test, data are reported as the mean (SD < 12% of each respective mean). **p<0.01, ***p<0.001, compared to ISC group [F (4, 27) = 15,84 p<0.0001].

#### Elevated Plus-maze Task (EPMT)

Figure 7. The elevated plus-maze task is widely used to study anxiety-like behavior. The task was performed on the 7^th^ day after treatment with EVs. ISC animals spent more time on the closed arms than the naive animals, indicating an anxiogenic-like effect of the stroke. This effect was reversed with EV1 or EV2 treatment.

**Figure 7.**
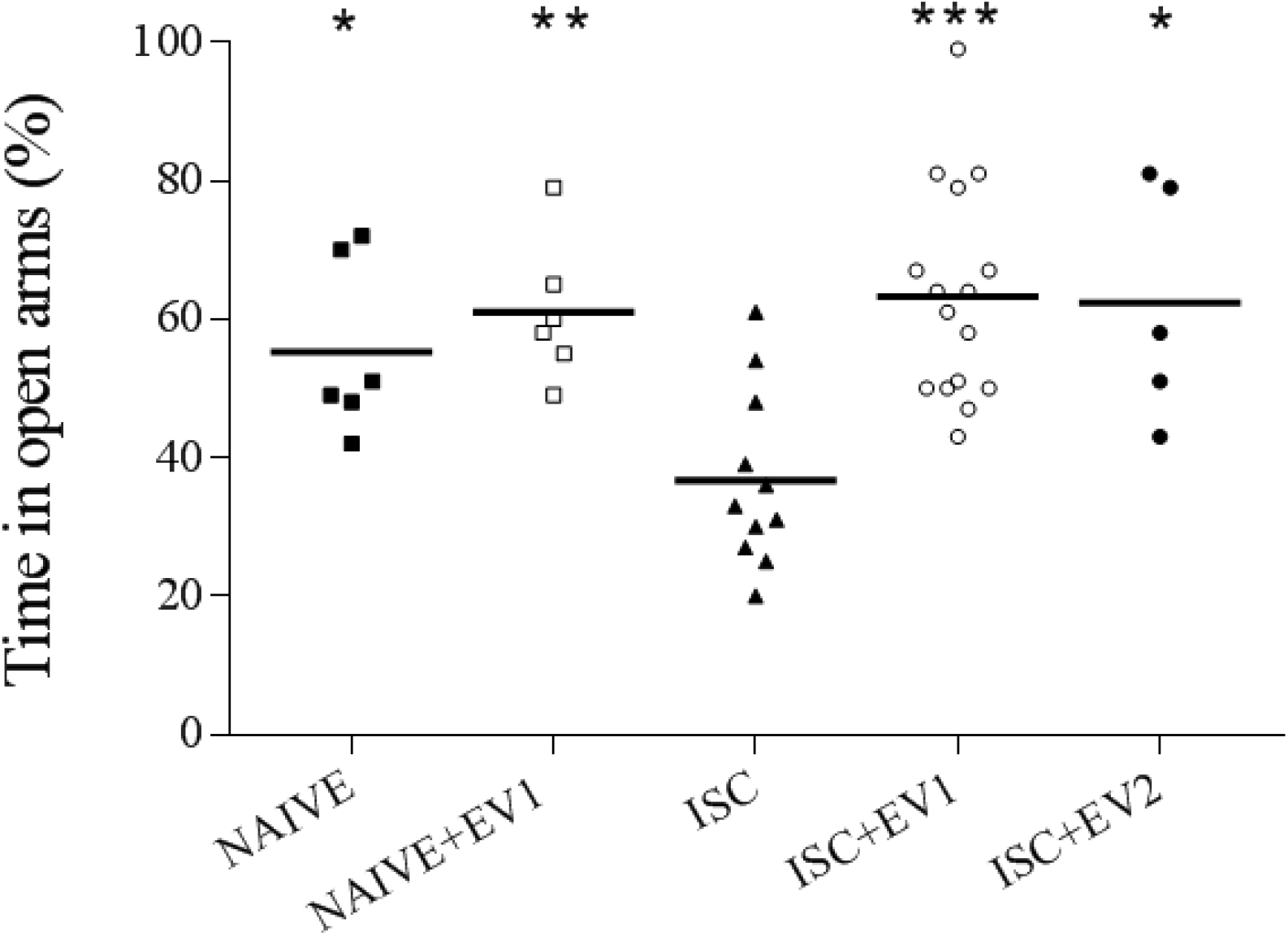
Elevated plus-maze. Groups: Naive (n=6), Naive+EV1 (n=6), ISC (n=11), ISC+EV1 (n=16) and ISC+EV2 (n=5). Data are reported as Mean (SD < 22% of each respective mean). 1-way ANOVA of a post-hoc Turkey’s test, *p<0.05, **p<0.01, ***p<0.001, compared to ISC group [F (4, 39) = 6,915 p=0.0003].

## DISCUSSION

In the present study, the focal permanent stroke model in rats, impaired: i) front paw symmetry; ii) working memory, short- and long-term memory; and iii) anxiety-like behavior. Interestingly, a single intranasal dose of EVs, administered 24 h after the insult, reversed these stroke sequels in a long-term manner. Additionally, EVs presented tropism to the penumbra zone, by comparing to other brain regions. In fact, it has been previously reported that EVs administration in rat models of brain diseases presents neuroprotective properties^31, 32, 50^ and tropism to the penumbra zone^35^. However, our study is the first demonstrating that a single intranasal dose of EVs, performed 24 h after a focal permanent stroke in rats, presented tropism to the penumbra zone and long-term neuroprotective activity. It is important to emphasize that all of these beneficial properties were induced by EVs released from hAT-MSCs, which could be easily obtained from a healthy individual.

Clinical data from stroke patients have shown that behavioral and motor impairment are dependent on the brain region where the core and the penumbra zone are developed^2, 5^. Our group had previously characterized, in a rat model of ischemic stroke, that the core and the penumbra zone are commonly detected in the prefrontal cortex and hippocampus^51^. As these brain regions are involved in memory modulation^52, 53^, we suggest that a single intranasal dose of hAT-MSCs-derived EVs may reach these regions and recover behavioral impairment by putatively reestablishing the blood flow. In fact, improvement of angiogenesis is currently acknowledged as an outcome of EVs transfer protein, mRNA and miRNA to endothelial cell^54^, regulating protein transduction^55, 56^. Additionally, clinical evidence demonstrated that depression and anxious behavior are outcomes of ischemic stroke^57, 58^. Accordingly, here, we observed that anxiety-like behavior caused by stroke was reverted by hAT-MSCs EVs.

Concerning motor disabilities, contralateral upper limb paresis is also a consequence of ischemic stroke in humans^7^. This outcome is well characterized by asymmetry of front paws in ischemic animals^40, 59^. Indeed, we observed that intranasal EVs treatment fully reversed this asymmetry of front paws caused by stroke.

These results shed light into a new and straightforward therapeutic strategy for stroke treatment, by utilizing the human adipose tissue as an accessible source for hAT-MSCs. The intranasal EVs administration after onset caused a long-term neuroprotection, offering a broader therapeutic window, compared to current standard routes. Together, these findings show the potential of hAT-MSCs-derived EVs as therapeutic agents in human patients with ischemic stroke.

## Authors’ contributions

Francieli Rohden, is responsible for the concept and design, participated in data acquisition, performed statistical analysis, drafting and revising the manuscript.

Luciele Varaschini Teixeira, Luis Pedro Bernardi and Nicolly Paz Ferreira Marques, participated in data acquisition, performed statistical analysis, drafting and revising the manuscript.

Mariana Colombo participated by analyzing vesicle size and purity and revising the manuscript.

Geciele Rodrigues Teixeira, Fernanda dos Santos de Oliveira, Elizabeth Obino Cirne Lima participated by isolating patient cells, discussing concepts and results, and revising the manuscript.

Fátima Costa Rodrigues Guma participated in the study concept and results and revising the manuscript.

Diogo Onofre Souza, was responsible for the concept and design of the study, obtaining funding and revising the manuscript.

