## Supplementary material for "Extracellular vesicles from human adipose stem cells are neuroprotective after stroke in rats": Supplemental material.docx

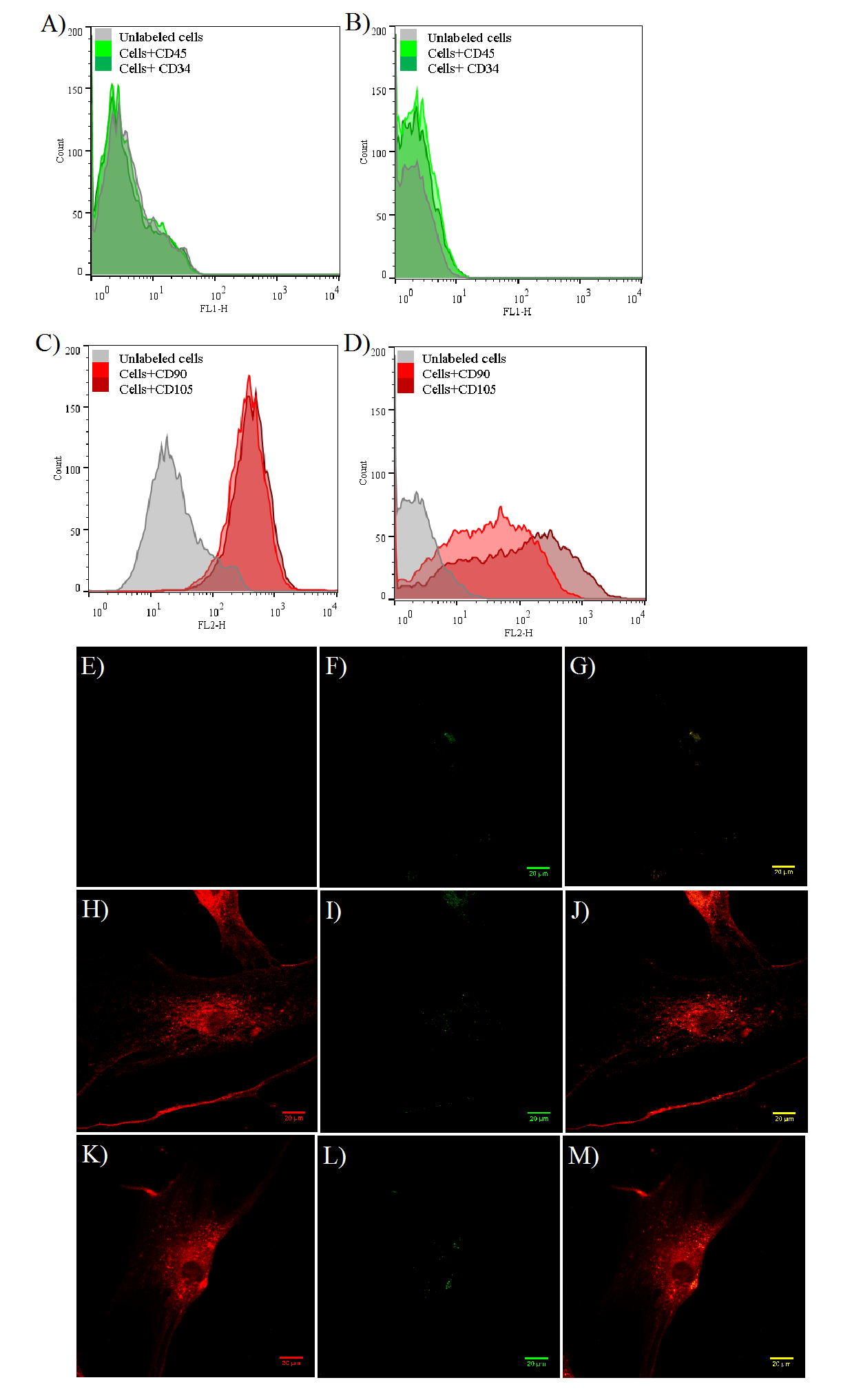
**Supplementary figure 1.** hADSCs cells characterization, labeling for specific proteins (clusters of differentiation) CD45, CD34, CD90 and CD105.

**Figure 1:** Representative images of C1 (A and C) and C2 (B and D) cells by flow cytometry and fluorescence microscopy with CD labeling specific for hADSCs (E-M). **A) and B)** CD45 and CD34 (non-expression in both cells). **C) and D)** CD90 and CD 105 (marking 70% of the cells). **E) to M)** Representative images of cells from fluorescence microscopy, 40x objective: **E)** Negative control cells for antibody Alexa fluor 555 anti-mouse (red); **F)** Negative control cells for antibody Alexa fluor 488 anti-mouse (green); **G)** Negative control cells for both antibodies merge: Alexa fluor 555 and for Alexa fluor 488; **H)** CD90 (red); **I)** CD45 (green); **J)** Merge for CD90 and CD45; **K)** CD105 (red); **L)** CD34 (green); **M)** Merge for CD105 and CD34. Scale bars: 20µm

**Supplementary figure 2.** Symmetric score of animals evaluated through Cylinder Task (CT).

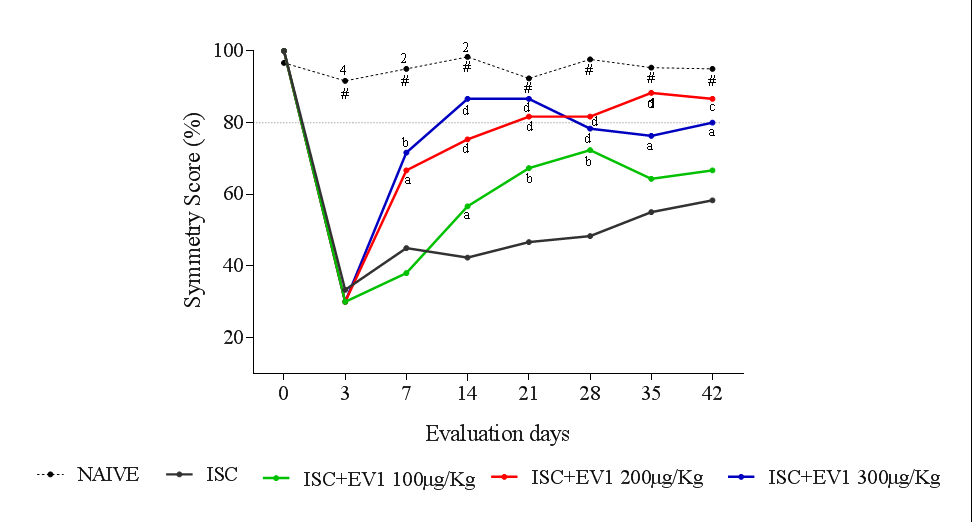

**Figure 2.** Dose curve for: Naive, ISC, ISC+ EV1 100μg/kg, ISC+EV1 200μg/kg, ISC+ EV1 300μg/kg, n = 3 for each group. Day 0 refers to baseline symmetry, evaluated 24 hours before induction of stroke. Data are expressed as mean (SD were less than 32% of respective mean) and analyzed by two-way ANOVA followed by Tukey’s test: ^a^p < 0.05, ^b^p < 0.01, ^c^p < 0.001 and ^d^p < 0.0001, compared to the ISC group; ^2^p < 0.01, ^4^p < 0.0001, compared to 200µg/Kg and 300µg/Kg group. ^#^p<0.01 compared to 100µg/Kg.
